# ESKtides: a comprehensive database and mining method for ESKAPE-derived peptides

**DOI:** 10.1101/2023.05.03.538019

**Authors:** David Runze Li, Hongfang Wu, Geng Zou, Xuejian Li, Yue Zhang, Yang Zhou, Huanchun Chen, Jinquan Li

**Affiliations:** State Key Laboratory of Agricultural Microbiology, Key Laboratory of Environment Correlative Dietology, College of Food Science and Technology, Shenzhen Institute of Nutrition and Health, Huazhong Agricultural University, Wuhan, 430070, China; Hubei Hongshan Laboratory, College of Biomedicine and Health, Huazhong Agricultural University, Wuhan, 430070, China; Shenzhen Branch, Guangdong Laboratory for Lingnan Modern Agriculture, Genome Analysis Laboratory of the Ministry of Agriculture and Rural Affairs, Agricultural Genomics Institute at Shenzhen, Chinese Academy of Agricultural Sciences, Shenzhen, 518000, China

**Author notes:** Correspondence (Jinquan Li). These authors contributed equally: David Runze Li, Hongfang Wu, Geng Zou.

## Abstract

Under the concept of one health, Super-drug resistant bacteria are getting more and more attention from scientists, in particular, ESKAPE bacteria directly killed more than 1,270,000 people in 2019. Recent studies have proposed that phage PGHs(peptidoglycan hydrolases) and antimicrobial peptides can be the new antibacterial agents against multi-drug resistant bacteria. However, Methods for mining antimicrobial peptides based on phages or phages PGHs are in need to develop. Here, using ESKAPE strains and ESKAPE phages in total of 6,809 samples from National Center for Biotechnology Information (NCBI), PhagesDB, Microbe Versus Phage (MVP) and Virus-Host Database, we systematically identified PGHs across ESKAPE strains prophages and phages, mined peptides and scored peptides based on PGHs. As a result, a total of 1,000 high antibacterial activity peptides and 1,200 medium antibacterial activity peptides were identified. In order to reduce the impact of different methods on comments, we used a unified process to comment ESKAPE strains and phages. In addition, we designed an online tool to predict the peptides antibacterial activity. Calculations of peptides phylogeny, peptides physicochemical property and peptides secondary structure are also included into our online tools. Finally, we developed ESKtides, a user-friendly and intuitive database (http://www.phageonehealth.cn:9000/ESKtides) for data browsing, searching, and downloading. ESKtides will significantly provide a rich peptide library based on ESKAPE strains and phages.

## INTRODUCTION

ESKAPE bacteria drug resistance is a serious medical problem. Into the 21st century, scientists in multiple fields are actively promoting global health. In recent years, the concept of one health is proposed and always reminds us that human health is not isolated but connected to the health of animals, plants and microorganism[1]. Therefore, the topic of one health shows an increasing trend from 2000-2020, especially after the conference on zoonotic diseases to environmental degradation, human and animal pathogens were focused on [2]. In 2019, roughly 1,270,000 peoples died due to global AMR infections, by 2050, the AMR will cause 10 million deaths each year and the total direct economic loss of 3 trillion pounds [31]. Under the ‘One health’ frame, some representative strains that pose a serious threat to ‘ One health’ is ESKAPE (*Enterococcus faecium, Staphylococcus aureus, Klebsiella pneumoniae, Acinetobacter baumannii, Pseudomonas aeruginosa* and *Enterobacter* species). As early as 2017, WHO published a list of pathogens and ESKAPE is listed as priority status [3]. Currently, incorrect use of broad-spectrum antibiotics has led to an increasing number of pathogenic bacteria developing resistance to the many antibiotics. In this field, we can not ignore an important problem: antimicrobial resistance. Until now, AMR infections evens still linger and caused huge damage of life and property. Up to now, researchers concluded ESKAPE strains drug-resistant spectrum: oxazolidinones, lipopeptides, macrolides, fluoroquinolones, tetracyclines, beta-lactams, beta-lactam-beta-lactamase inhibitor combinations[5], Obviously, these drug-resistant spectrum almost covers more than 90% of the types of antibiotics. Therefore, traditional antibiotic therapy has been unable to face with the increasing problem of drug resistance, since 2017, many therapeutic schedules were proposed by U.S. Food and Drug Administration (U.S. FDA)[8], which are mainly divided into two aspects: 1. explore new antibacterial targets; 2. develop new antibiotics.

Phage, phage PGHs(peptidoglycan hydrolases) and antimicrobial peptides are the new antibacterial agents against multi-drug resistant bacteria. At present, many new antibacterial agents are developed in order to use against multi-drug resistance, especially for ESKAPE bacteria. As a new antibiotic substitute, the source of the antimicrobial peptides are widely distributed, meanwhile, antimicrobial peptides are the small molecular proteins that it’s more likely to be absorbed by the body, shorter and more easily accessible antimicrobial peptides are considered as candidates in clinical translation. Most antimicrobial peptides are positively charged cationic peptides and amphiphilic. According to the net charge of the amino acid sequence, protein structure and the source, peptides are divided into several subsets: 1. Anion antimicrobial peptides; 2. Cation alpha-helix antimicrobial peptides; 3. Cation beta-sheet antimicrobial peptides; 4. Cationic antimicrobial peptides containing special amino acids; 5. Peptide fragments of antimicrobial proteins[9]. Until now, bactericidal mechanism of antibacterial peptides are divided into two aspects: 1. the cell membrane targeting mechanism that uses hydrophobic action to destroy cell membranes and form holes to cause cell death[10]; 2. Intracellular targeting mechanisms that destroy cells by interfering with their normal metabolism[11,12,13]. According to related studies, most natural antimicrobial peptides come from animals, plants, bacteria, fungi and protozoa. As a kind of virus, phages are widely distributed in the environment and in large quantity (according to statistics, there are 1031 phages in nature). Phages as bacterial hunters can specifically lyse bacteria, meanwhile phage-encoded proteins such as phage PGHs perform antibacterial activity and phage PGHs will cleave the host internally to assist release[15-17]. Thus phages contain huge potential for antimicrobial peptides mining. With the advent of the era of high-throughput sequencing and culturomics, more and more phages and prophages are being discovered. However, large-scale genome-wide analyses of phage or prophage antimicrobial peptides have rarely been reported, and no database for antimicrobial peptides in phages and prophages are available, especially compared with DRAMP 2.0 [26] DRAMP 3.0 [27] and APD3 [28]. Recently, Thandar et al analyzed the secondary structure of phage lysin(a kind of PGHs) PlyF307 and intercepted the C-terminal antimicrobial peptide P307 containing double alpha-helical structure and also verified bactericidal activity in vitro[14], which imply we can mine peptides by PGHs, two independent studies indicate that people also can use deeplearning model to predict antimicrobial peptides activity[1,14]. Therefore, it is feasible to add phage or prophage antimicrobial peptides based on PGHs mining as an additional dimension to the existing antimicrobial peptides dataset and use deeplearning model scoring.

In this study, by using ESKAPE strains genome data and corresponding phages genome data, we developed a new computational pipeline to systematically perform mining peptides analyses based on PGHs (peptidoglycan hydrolases) across ESKAPE strains and phages (for ESKAPE-derived peptides, https://github.com/hzaurzli/phatides_prediction). We further scored ESKAPE-derived peptides antimicrobial activity, calculated peptides physiochemical properties and analysed secondary structure of peptides. The ESKtides database (http://www.phageonehealth.cn:9000/ESKtides) was constructed for browsing, searching, analyzing the ESKAPE-derived peptides data.

## MATERIALS AND METHODS

### Collection and processing of data for ESKAPE strains and phages

We downloaded genome data (including ESKAPE strains and their phages) from four widely used databases on January 2023, including MVP [31], PhagesDB [32], VHDB [33] and NCBI [34], and merge these genomes to construct our dataset in this study. The data in these four databases were subjected to the following processing. First, for ESKAPE strains, we filtered incomplete assemblies (such as contigs, scaffolds) to ensure that the accuracy of mining. Second, for ESKAPE phages, we downloaded the phages genome corresponding to the their strains (phages are isolated from corresponding strains) from MVP, PhagesDB, VHDB and marked as corresponding strains’ phages. Removing genomes include phages that do not belong to ESKAPE strains or prophages genome from metagenome. Third, we used checkM [29] and checkV [30] to evaluate genome integrity and filter the data which integrity is less than 90%. In order to avoid artificial differences resulting from different annotation pipelines, in this study, all the assemblies or genomes were reannotated with prokka. The genome annotation circos was generated by CGView.js.

### ESKAPE-derived peptides mining

In this study, different analysis pipelines have been developed for ESKAPE strains and phages. For strains, each strain ORFs were done by using prokka 1.14.6 [18]. Phispy 4.2.21 [19] used to discover prophages based on annotations and then prophages coordinate was extracted by custom scripts. To re-annotate prophages genome, phanotate[20] was performed to annotate prophages ORFs and protein sequences were obteined from ORFs. CDhit [21] was used to eliminate redundancy and remove proteins which molecular weight is more than 40 kDa. Enzymes were searched in CAZyme which cleat sugar chains as substrates by using ‘run_dbcan.py’[22]. Reported PGHs domains were included into our pipeline by using ‘hmmsearch’[23]. Next, Signaling proteins and transmembrane proteins are removed which predicted by SignalP v6.0 [24] and TMHMM 2.0. Finally, we obtain all putative PGHs. Peptides with a length of 6-50aa were obtained by using a sliding window based on PGHs sequences and the activity fraction was calculated. For phages, phanotate[20] was performed to annotate phages ORFs and protein sequences were obteined from ORFs directly. The remaining steps are the same as strains.

### Activity score prediction and secondary structure calculation

The basic model is a deep learning frame with convolutional NNM and LSTM layer, which has been used in the current research of Ma et al [1]. The model was constructed by following steps: 1. The non-AMP datasets were downloaded from UniProt (http://www.uniprot.org) and AMP data were mainly collected from four public AMP datasets (ADAM, APD, CAMP and LAMP). The whole dataset was first split into two sets at a ratio of 8:2 for training set and testing set; 2. The basic layers and parameters were set: the embedding layer, input dimension is 21, output dim is 128 and input length is 300; the 1D Convolution layer: number of convolution kernels is 64, length of each filter is 16, activation function is relu(); 1D max pooling layer: pool size is 5; LSTM layer: units is 100; and dense layer: units is 1, activation function is sigmoid. 3. Using training dataset to train models, the models were built with the Keras framework (version 2.2.4, https://www.keras.io). For secondary structure, we used SCRATCH-1D 1.1 [25] to calculate.

### Evaluate the physiochemical properties of the peptide

We also provide a platform for users to calculate physicochemical property by using R package ‘Peptides’ (https://www.rdocumentation.org/packages/Peptides/versions/2.4.4) and Biopython (version 1.79). Protein length, molecular weight, instability, hydrophobicity, hydrophobic moment, aliphatic, pI and charge were included in our platform by using different functions:‘lengthpep()’,‘mw()’,‘instaIndex()’,‘hydrophobicity()’,‘hmoment()’,‘aIndex()’, ‘pI()’ and ‘charge()’ in R package Peptides. Protein gravy is calculated by function ‘gravy()’ in Biopython.

## IMPLEMENTATION

The ESKtides website runs on an Apache Web server (https://apache.org/). The database was developed by using sqlite3 (https://www.sqlite.org/index.html). Flask 2.0.3 (https://flask.net.cn/) was used for server-side scripting. The ESKtides web interface was built by using Datatables (https://datatables.net/) and JQuery v2.1.1 (http://jquery.com). ECharts(http://echarts.baidu.com) was used as a graphical visualization framework and R (https://www.r-project.org/) for graph drawing. We recommend using the latest versions of Firefox and Google Chrome for the best experience.

## DATABASE CONTENT AND USAGE

### Samples in ESKtides

In total, 5,630 ESKAPE strains and 1,179 ESKAPE phages were included in ESKtides, For ESKAPE strains, our dataset is ranging from 1,132 samples in Staphylococcus aureus to 921 samples in Escherichia coli and for ESKAPE phages, which is ranging from 189 samples in Staphylococcus aureus phages to 418 samples in Escherichia coli phages (Table 1). The detailed information, including the number of samples per strains and phages, reference genome versions and the number of ORFs, is available on the ‘Browse Genome’ page. As a user-friendly data portal, ESKtides displays ESKAPE related information across a large number of strains and phages.

**Table 1.**
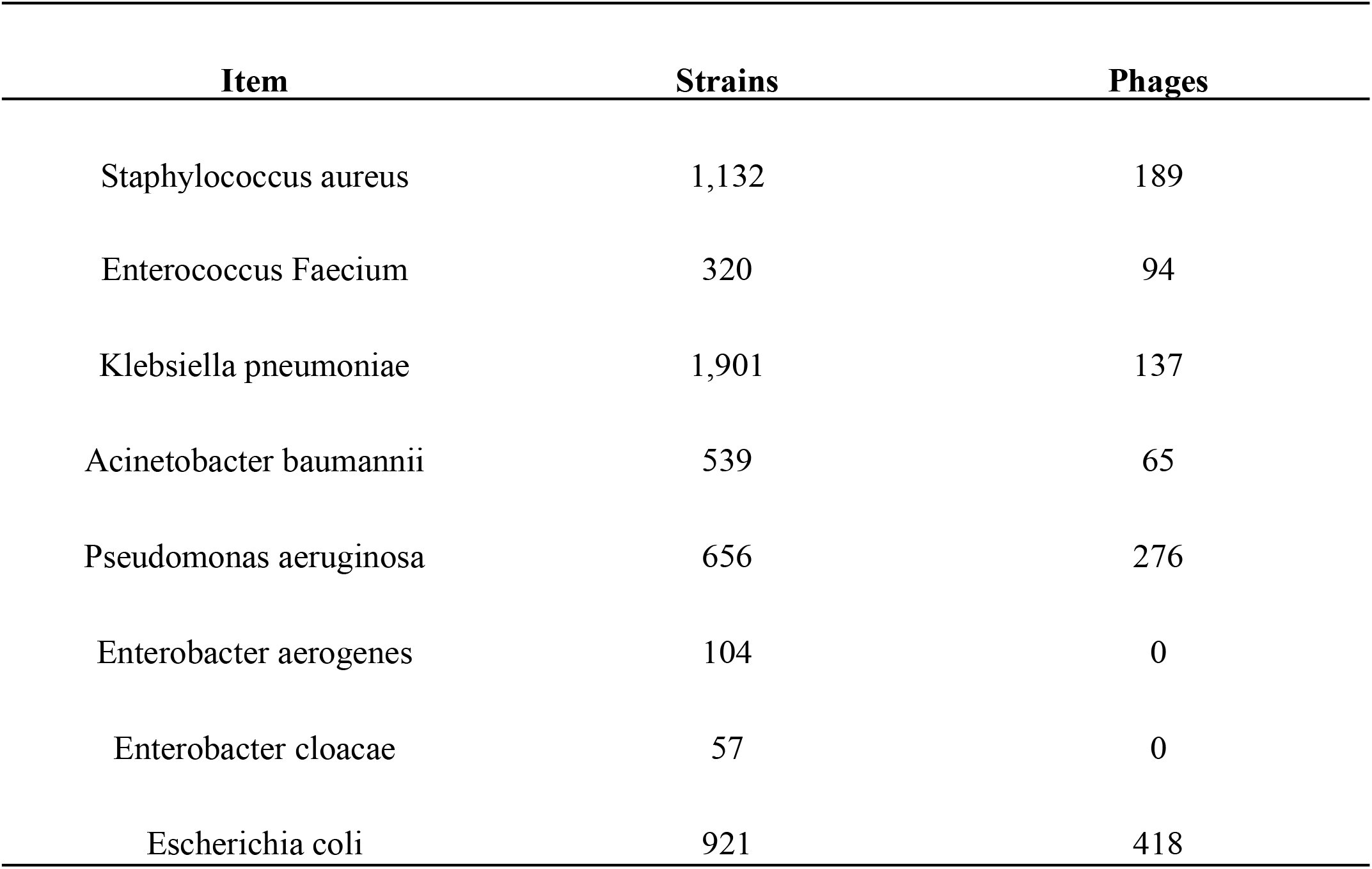
ESKAPE dataset

### ESKAPE-derived peptides distribution in ESKAPE strains and phages

Our approach is based on PGHs(peptidoglycan hydrolases) for peptides mining, We identified a total of 185,177 PGHs in these strains and phages, ranging from 1,132 in *Staphylococcus aureus* to 418 in *Escherichia coli* phages at different strains and phages. For ESKAPE-derived peptides, a total of 22,091,402 active peptides were found and 12,067,248 peptides reveal high activity (Table 2). Many ESKAPE-derived peptides are mined in multiple strains and phages, which provides a rich peptides library for subsequent research. In our database, different strains have different mining efficiency, *Enterobacter* (*Escherichia coli, Enterobacter cloacae* and *Enterobacter aerogenes*) with the maximum mining efficiency (*Escherichia coli*: 2921 per strain, *Enterobacter cloacae*: 24579 per strain, *Enterobacter aerogenes*: 5662 per strain), *Staphylococcus aureus* with the minimum efficiency(305 per strains). For phages, *Enterococcus Faecium* phages have the maximum efficiency(1315 per phage) and *Pseudomonas aeruginosa* phages have the minimum efficiency (850 per phage).

**Table 2.** ESKAPE-derived peptides

| Item | Strains derived peptides | Phages derived peptides |
| --- | --- | --- |
| Staphylococcus aureus | 345,260 | 256,255 |
| Enterococcus Faecium | 552,969 | 123,647 |
| Klebsiella pneumoniae | 2,199,679 | 116,563 |
| Acinetobacter baumannii | 1,100,829 | 101,998 |
| Pseudomonas aeruginosa | 902,600 | 234,596 |
| Enterobacter aerogenes | 588,854 | 0 |
| Enterobacter cloacae | 1,401,037 | 0 |
| Escherichia coli | 2,690,162 | 431,252 |

### Web interface

We designed a user-friendly web serve for the database, five main modules, including ‘Browse Genome’, ‘Browse Peptides’, ‘Analysis’, ‘Statistic’ and ‘Download’ (Figure 3A) are provided for users. In each table, search boxes are designed for users to quick search, with two patterns: 1. exact search; 2. fuzzy search. On the home page, ESKAPE-derived peptides pipeline was showed on the home page, user can better understand the whole mining process.

**Figure 1.**
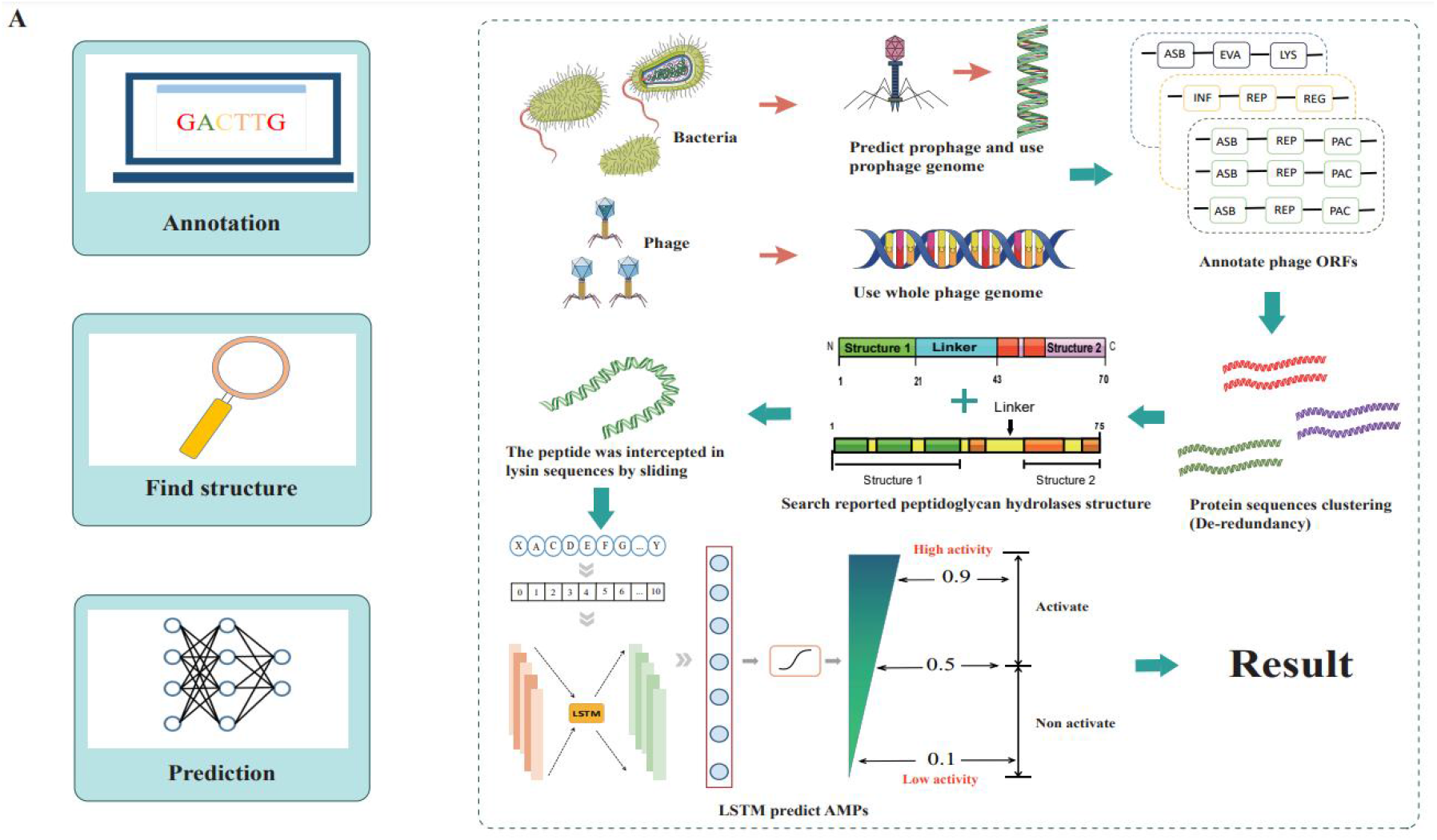
Worlflow for ESKAPE-derived peptides mining.

**Figure 2.**
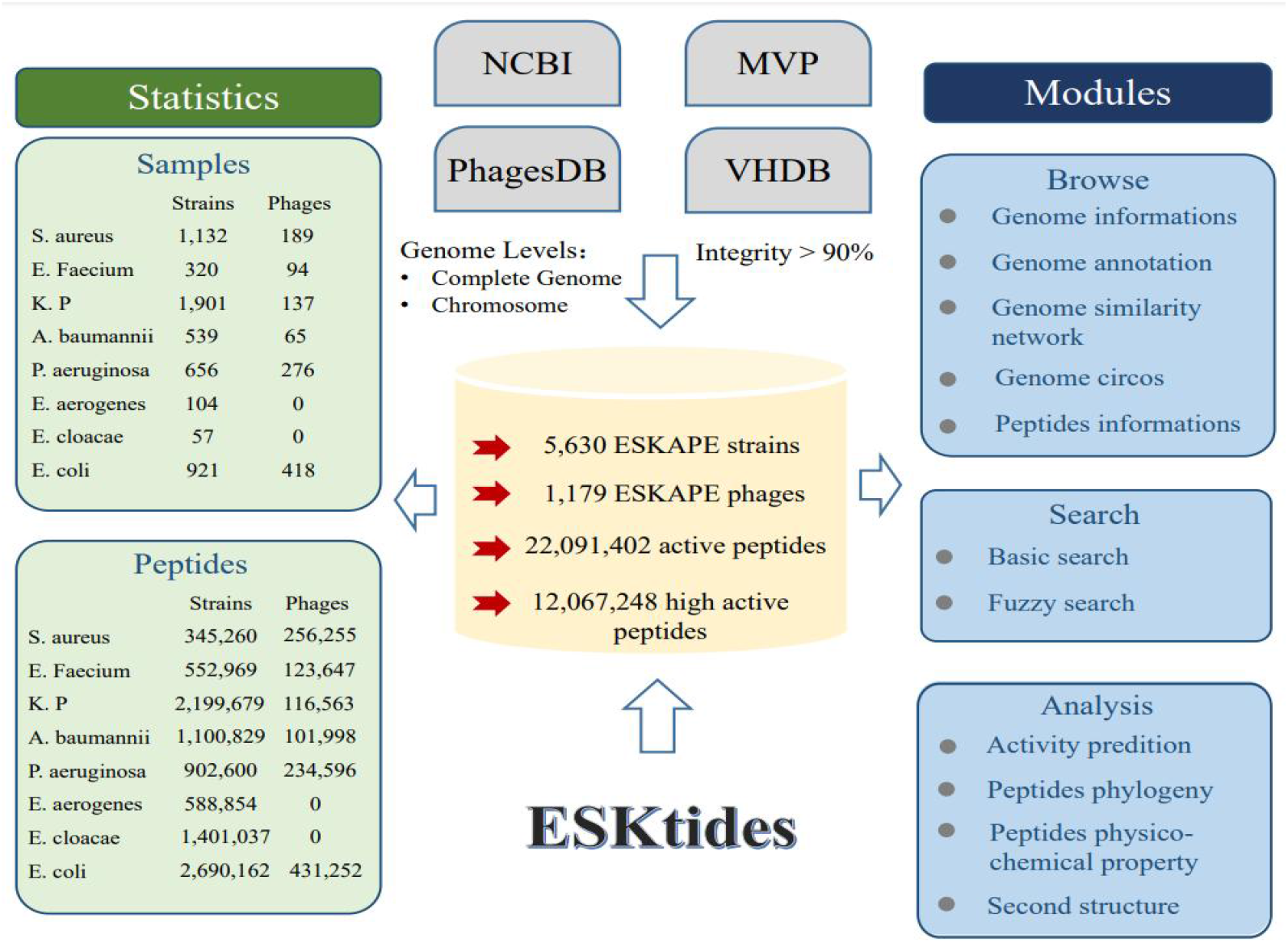
Overall design of ESKtides. ESKtides curated metadata information from NCBI, MVP, Virus-Host Database and phagedb, and the data in these four databases are subjected to the following processing. First, for ESKAPE strains, we filter incomplete assembly to ensure that the accuracy of mining. Second, for ESKAPE phages we downloaded the phages genome corresponding to their strains (phages are isolated from corresponding strains) from MVP, PhagesDB, VHDB and marked as corresponding strains’ phages. All raw data was processed by using a standard pipeline. ESKtides includes ‘Browse’, ‘Search’, ‘Download’ and ‘Submission’.

**Figure 3.**
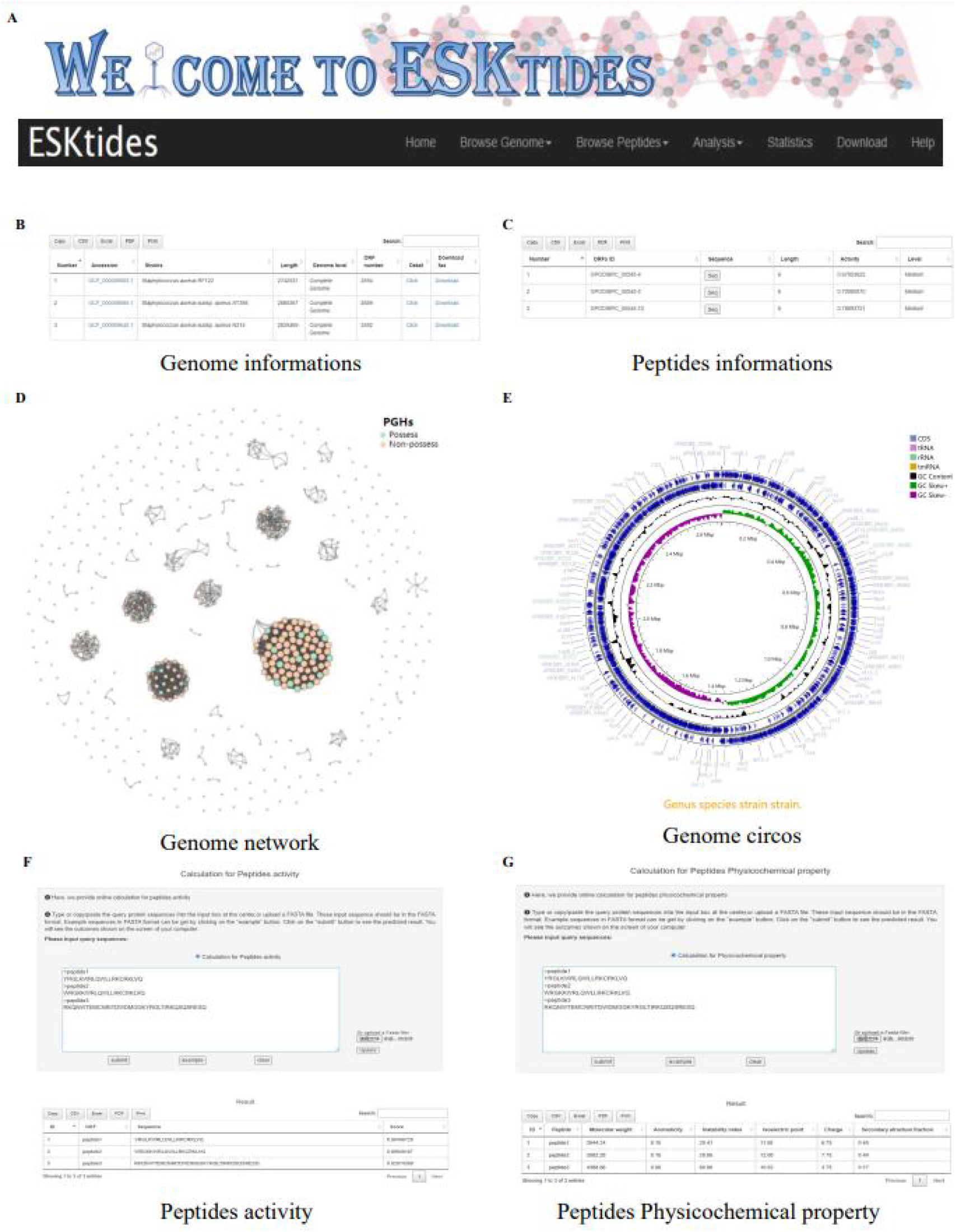
Overview of ESKtides. (A) Main functions of ESKtides, including the ‘BrowseGenome’,‘BrowsePeptides’,‘Analysis’, ‘Statistic’, ‘Download’ and ‘Help’ modules. (B) A table of queried strains or phages information in the ‘Browse Genome’ module. (C) A table of queried ESKAPE-derived peptides information in the ‘Browse Peptides’ module. (D) A graph of queried PGHs distribution in the ‘Browse Genome’ module. (E) The annotation circos graph of queried strain. (F) Submit peptides sequences to score the bactericidal activity of the peptide. (G) Calculate peptides sequences physicochemical property.

On the webpage of ‘Browse Genome’, users can browse ESKAPE profiles of different strains and phages on the top box, search genome informations by accession, strains, genome length, genome level and ORF number on the center table and browse genome similarity network on the bottom graph, each point represents one genome, we colored two groups to indicate PGHs distribution that one group is PGHs and another group is not PGHs. For example, if users want to query genome annotation, they only need to click button ‘detail’ to annotation page, which is on the center table. The searching results will be exhibited in a table that contains ‘Locus tag’, sequence type, ‘CDS length’, ‘gene name’, ‘EC number’, ‘COG id’ and ‘product function’ (Figure 3B). In addition, users can browse genome annotation circos on the top and also can download annotated proteins fasta files by clicking ‘Download’.

On the webpage of ‘Browse Peptides’, users can view the ESKAPE-derived peptides from different ESKAPE strains or phages and also can search peptides informations by ‘ORFs ID’, ‘Sequence’, ‘peptides length’, ‘peptides activity’ and ‘activity level’. The antibacterial activity value is within 0-1. Two levels are set for activity level, if peptides activity is more than 0.9, which is defined ‘High’, if peptides activity is within 0.5-0.9, defined as ‘Medium’, otherwise we do not consider inclusion in our dataset. Users can select ESKAPE-derived peptides of interest and click bottom ‘Seq’ to download (Figure 3C). Group information contains ‘ORFs ID’, ‘Sequence’, peptides length, peptides activity and activity level, users also can download all the ESKAPE-derived peptides on the ‘Download’ page in corresponding group datasets that users are interested in.

On the webpage of ‘Analysis’, in ‘Peptides activity Prediction’, users can predict bactericidal activity of peptides by using deep learning model. Users can start a new calculation after clicking the ‘submit’ button and click ‘example’ button to show example format, ‘clear’ button can erase the previous record. In ‘Peptides Phylogenetic tree’, users can show the phylogenetic relationship of related peptides, user only need to upload tree file (newick format or json format) and adapt different styles to your needs. We also provide physicochemical property calculation platform for peptides, which is used in the same way as ‘Peptides activity Prediction’. We also provide the ability to predict secondary structure in ‘Peptides secondary structure’ by using SCRATCH-1D_1.1[31], all operations can be performed in batches.

### Download

All the analysis results can be downloaded as CSV files of each group on the ‘Download’ page and all the search and calculate results can be downloaded as CSV and Excel files for customized analysis by clicking the corresponding button on the top right of almost all tables.

### Data submission

Users can submit relevant data by sending us a data information table via email. Currently, ESKtides can accept open access genomes fasta files and assemblies fasta files, which are related ESKAPE. The submitted data would be added to ESKtides after curation and analysis as described in the section of Materials and Methods. We also provide ESKAPE-derived peptides mining pipeline, The software is available on GitHub: https://github.com/hzaurzli/phatides_prediction.

### Case study

In this section, we conduct a case study to estimate the performance of our activity prediction platform and then predict reported peptides based on PGHs. We first searched all the relevant papers which are confirmed by biological experiments and recorded all peptides sequences before 1 January 2023. Then, we ranked the peptides activity according to the prediction score and searched the newly published literatures to verify whether the predicted peptides have been confirmed by biological experiments. We list these predicted peptides activity in Table 3 (sorted by prediction scores). Compared with CAMPR4, our platform (deep learning model in Ma et al [1]) has more advantages in predicting performance of peptides mining based on PGHs. For example, the analysis results in papers[2,5] described that *Pseudomonas aeruginosa* peptides (X1-X4 and PaP1-1 from PGHs) show low bactericidal activity in experiment, but CAMPR4 was determined to be high activity, our platform was determined the low activity, which is more fitted with experiment result. The results of this case study demonstrate that our platform do better in identifying peptides and narrowing the scope of candidates for further biological experiments.

**Table 3.** Case study

| Source | Peptides | Experimental | Activity score | Activity score | Liter |
| --- | --- | --- | --- | --- | --- |
| Pseudomonas aeruginosa PGHs | X1 | Low | 0.029 | 0.81(RF) | [5] |
| Pseudomonas aeruginosa PGHs | X2 | Low | 0.016 | 0.85(RF) | [5] |
| Pseudomonas aeruginosa PGHs | X3 | Low | 0.016 | 0.84(RF) | [5] |
| Pseudomonas aeruginosa PGHs | X4 | Low | 0.26 | 0.53(RF) | [5] |
| Pseudomonas aeruginosa PGHs | PaP1-1 | Low | 0.382 | 0.64(RF) | [2] |
| Pseudomonas aeruginosa PGHs | PaP1-2 | Low | 0.757 | 0.82(RF) | [2] |
| Acinetobacter baumannii PGHs | P307 | Good | 0.999 | 0.94(RF) | [6] |
| Acinetobacter baumannii PGHs | P103 | Low | 0.005 | 0.29(RF) | [3] |
| Acinetobacter baumannii PGHs | P104 | Good | 0.029 | 0.06(RF) | [3] |
| Mycobacterium PGHs | AK15 | Excellent | 0.937 | 0.97(RF) | [4] |

## SUMMARY AND FUTURE DIRECTIONS

Recent advantages in experimental methods and available sequencing techniques have led to an explosive growth of biological data, especially in microorganism and viruses. It is worthwhile to mine peptides with bactericidal activity in six super bacterias and their phages (ESKAPE). Many peptides-related databases such as DRAMP 2.0 [26] DRAMP 3.0 [27] and APD3 [28] have been constructed. However, there has been limited research on mining peptides by using pathogenic bacteria’s peptidoglycan hydrolase. In this study, we developed ESKtides by collecting public ESKAPE strains and phages genome data, which provides comprehensive information of ESKAPE-derived peptides in different ESKAPE strains and phages. To the best of our knowledge, ESKtides is the first and most comprehensive ESKAPE derived peptides database, we also provide some useful tools such as ‘Peptides activity Prediction’, ‘Peptides Phylogenetic tree’ and physicochemical property calculation to help users select more suitable peptides. In this version of ESKtides, by collecting the data of 5,630 ESKAPE strains and 1,179 ESKAPE phages. we systematically re-annotated the ESKAPE strains and phages, and identified PGHs in all kinds of ESKAPE strains and phages, obtained different length peptides based on PGHs and predicted activity of those peptides, which are expected to greatly expand our knowledge of peptides source and PGHs’ distribution in ESKAPE strains and phages. With the rapid development of artificial intelligence technology (AI), the performance and accuracy of predictions are also improving rapidly (AUC: greater than 92%) [1]. However, considering that only use AI to predict activity were caused false positives, the database probably incorporate some ESKAPE-derived peptides that are less active in the real situation. In the future, we will further do more confirmatory experiments to reduce false positive rate. With a comprehensive ESKAPE-derived peptides database in different ESKAPE strains and phages, we believe that ESKtides will be a valuable resource for understanding the functions and technology of drug synthesis of peptides.

## DATA AVAILABILITY

ESKtides is freely available to the public without registration or login requirements (http://www.phageonehealth.cn:9000/ESKtides).

## AUTHOR CONTRIBUTIONS

**David Runze Li**: Data curation, Methodology, Software,Writing – review & editing. **Hongfang Wu**: Data curation, Writing– review & editing. **Geng Zou**: Data curation, Methodology, Writing. **Xuejian Li**: Data curation, Formal analysis, Resources, Investigation. **Yue Zhang**: Resources, Investigation. **Yang Zhou**: Funding acquisition, Writing–review & editing. **Huanchun Chen**: Writing–review & editing. **Jinquan Li**: Conceptualization, Funding acquisition, Investigation, Resources, Software, Supervision, Validation, Visualization, Writing– review & editing.

## SUPPORTING INFORMATION

The online version contains supplementary figures and tables available.

